# The Hi-Culfite assay reveals relationships between chromatin contacts and DNA methylation state

**DOI:** 10.1101/481283

**Authors:** Elena K. Stamenova, Neva C. Durand, Olga Dudchenko, Muhammad S. Shamim, Su-Chen Huang, Yiqun Jiang, Ivan D. Bochkov, Suhas S. P. Rao, Eric S. Lander, Andreas Gnirke, Erez Lieberman Aiden

## Abstract

Hi-Culfite, a protocol combining Hi-C and whole-genome bisulfite sequencing (WGBS), determines chromatin contacts and DNA methylation simultaneously. Hi-Culfite also reveals relationships that cannot be seen when the two assays are performed separately. For instance, we show that loci associated with open chromatin exhibit context-sensitive methylation: when their spatial neighbors lie in closed chromatin, they are much more likely to be methylated.

## Main Text

Nuclear architecture and DNA methylation play critical and interdependent roles in mammalian genome regulation^1,2^. For example, the insulator protein CTCF, which interacts with the cohesin complex to form chromatin loops and thereby establish discrete structural and functional segments of the genome, binds in a methylation-sensitive fashion^3,4^. Similarly, a recent study has implicated hyper-methylation induced disruption of chromosome topology in oncogene activation^5^.

Nuclear architecture and DNA methylation are typically interrogated independently using Hi-C proximity ligation mapping^6^ and whole-genome bisulfite sequencing (WGBS)^7^, respectively. However, independent assays of architecture and methylation can obscure the interdependence of the underlying phenomena. In addition, since both sequencing assays require deep coverage of the genome, performing them separately increases the cost.

We have developed Hi-Culfite, a combined protocol integrating *in situ* Hi-C^8^ and bisulfite conversion^9^. We show that Hi-Culfite generates chromatin contact and DNA methylation maps whose quality and resolution rival those obtained by the respective single assays. Hi-Culfite data sets also allow integrated, multi-omics analyses that reveal unique biological insights, such as relationships between DNA methylation and spatial context, which cannot be obtained from separate Hi-C and WGBS data sets.

Hi-Culfite library construction (**Fig. 1A**) follows the initial steps of our *in situ* Hi-C protocol^8^: one million cells are crosslinked with formaldehyde; chromatin is digested with a restriction enzyme; and the overhangs are filled in, incorporating a biotinylated nucleotide to mark the fragment ends after ligation. The DNA is then purified and sheared. Next, we perform bisulfite conversion, which converts unmethylated cytosine residues into uracil but leaves methylated cytosine residues intact. We capture the resulting single-stranded, biotin-marked ligation products on streptavidin beads. Finally, we add adapters to the 3’ end, synthesize the 2nd strand by primer extension, ligate adapters to the 3’ end of the 2^nd^ strand, and PCR amplify with indexed primers (**Online Methods**).

**Figure 1.**
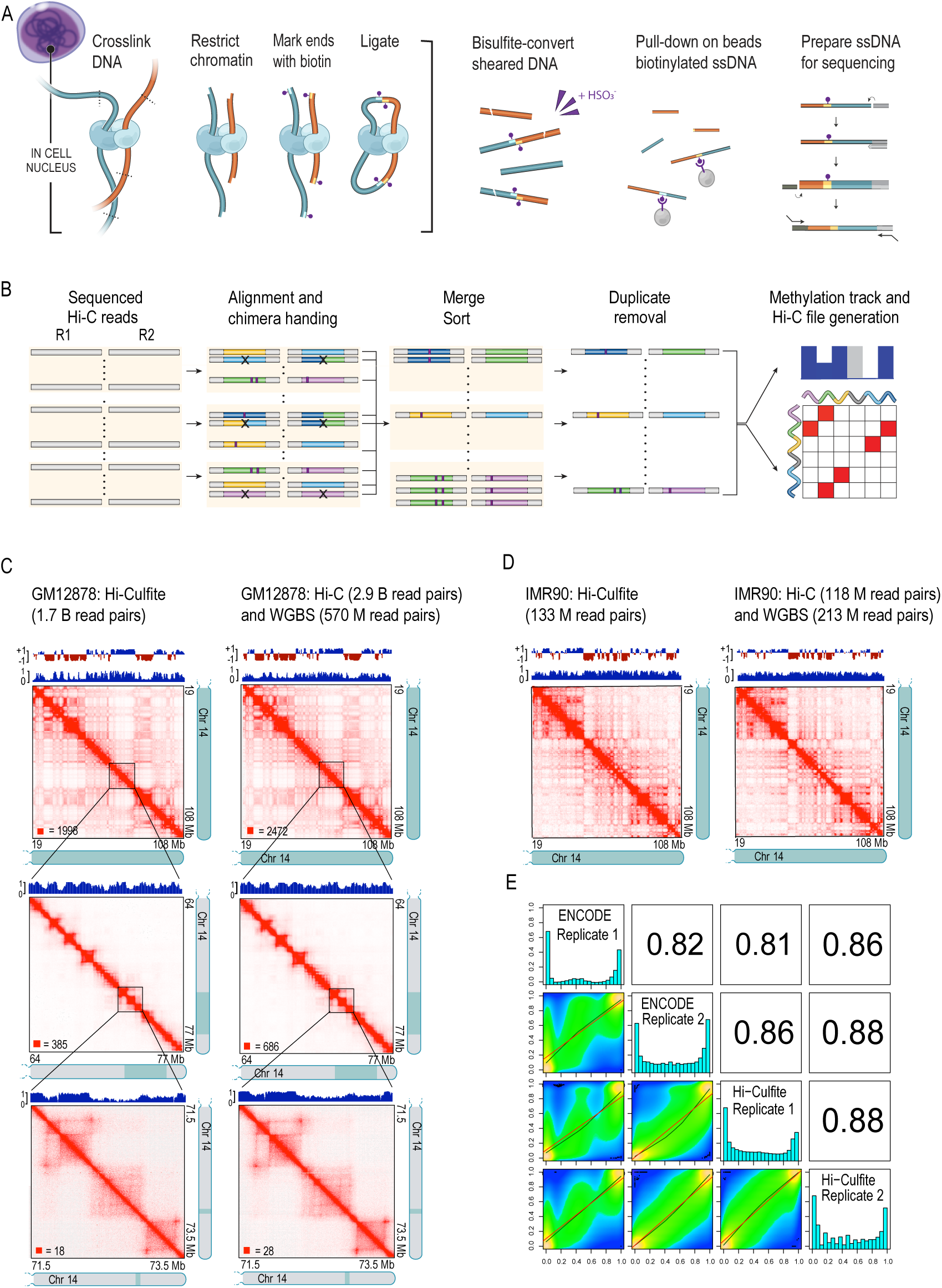
Hi-Culfite maps chromatin contacts and DNA methylation simultaneously. (**A**) Left: the initial steps are identical to standard *in situ* Hi-C. Right: the “bisulfite first” protocol entails capture of biotin-marked bisulfite-converted ligation products and the use of a commercial kit for preparing sequencing libraries from bisulfite-converted DNA. (**B**) Overview of the JuiceMe pipeline for processing of Hi-Culfite sequencing data. The software is built off of Juicer^20^; reads are aligned as single end via *bwa-meth*^22^, chimeric reads are handled by either pairing appropriately or removal, reads are sorted and merged, and duplicates and near duplicates are removed. Simultaneously, all reads that map are merged into a bam, duplicates marked, and the software *MethylDackyl* (https://github.com/dpryan79/MethylDackel) is used to create the CpG methylation track. (**C,D**) Comparison of GM12878 (**C**) and IMR90 (**D**) chromatin contact maps and DNA methylation tracks of human chromosome 14q obtained in a single Hi-Culfite experiment (left) or by separate in situ Hi-C and WGBS assays (right). Chromosome-scale contact matrices have 1 Mb resolution. Zoom-ins have 50 kb and 5 kb resolution. CpG methylation tracks (**Online Methods**) are shown above each Hi-C contact matrix. Eigenvectors denoting open (positive values; blue) and closed chromatin compartments (negative; red) along the chromosome are shown at the top. (**E**) Smoothed scatter plots (bottom left), Pearson correlation coefficients for pair-wise comparisons (top right) and bimodal distributions of single-CpG DNA methylation values (diagonal). ENCODE rep1 and rep2 are biological WGBS replicates. The Hi-Culfite data were from combined 2 technical replicates prepared from the same batch of cross-linked GM12878 cells (Hi-Culfite Replicate 1) and one biological replicate derived from different expansion of the same cell line (Hi-Culfite Replicate 2). The scatter plot is produced by *MethylKit*^23^; the red line is the linear regression fit and the green line is the polynomial regression fit.

Notably, we found that by performing bisulfite conversion *prior* to biotin pulldown and adapter ligation, we achieve much higher yield. By contrast, early iterations of the protocol, in which bisulfite conversion was performed on un-amplified Hi-C libraries with methylated adapters, led to great drops in library complexity (i.e., the number of unique sequencing templates in the library). This is presumably due to extensive strand breakage on account of the harsh chemical conditions during the bisulfite conversion leading to a reduction of the number of PCR-amplifiable molecules^10^. For instance, at the final amplification step, 10 cycles of PCR resulted in Hi-Culfite libraries with a mean concentration of 32nM, comparable to our standard *in situ* Hi-C protocol^8^. By contrast, the “adapter first” workflow produced ∼100 times less library (0.2 nM) despite starting with 10-fold more cells (10 million) and two additional PCR cycles.

A Hi-Culfite map comprises pairs of neighboring bisulfite-converted DNA sequence reads, each indicating the methylation state of two loci that might lie far apart along the genome, but that were spatially adjacent at the time of the assay. By quantifying the frequency with which pairs of loci are found adjacent, a Hi-Culfite map – like an ordinary Hi-C map – can be used to create a contact matrix showing the frequency at which pairs of loci co-localize. By quantifying how often loci are methylated, the Hi-Culfite map can be used to create a genome-wide methylation profile.

We sought to validate the quality of Hi-Culfite results by comparing contact matrices and methylation profiles generated using Hi-Culfite to those produced when *in situ* Hi-C and WGBS experiments are performed separately. To do so, we performed a Hi-Culfite experiment in GM12878 lymphoblastoid cells, for which both Hi-C^8^ and WGBS data sets (DCC accession: ENCSR890UQO^11,12^) are publicly available. After initial quality control of the Hi-Culfite libraries, we generated a deep Hi-Culfite dataset containing a total of 1.75 billion read pairs (**Supplementary Table 1**), comparable to the ENCODE standard for *in situ* Hi-C (2 billion).

We found that the chromatin contact maps produced by Hi-Culfite are comparable to *in situ* Hi-C maps across all resolutions we examined (1Mb → 5kb; **Fig. 1C**). Crucially, compartments (which arise when open [“compartment A”] and closed [“B”] chromatin segregate in the nucleus, manifesting as a plaid pattern in the contact map), contact domains (intervals in which all loci exhibit an enhanced contact frequency within themselves, manifesting as a bright square along the diagonal of the contact map), and loops (pairs of loci exhibiting an enhanced contact frequency with one another relative to their 1D genomic neighborhood, manifesting as peaks in the contact map) are evident in the Hi-Culfite data. Similarly, when we compared our results to ENCODE WGBS data, we found that the methylation profiles closely resembled one another at all resolutions. Indeed, the average single-CpG methylation correlation between Hi-Culfite and ENCODE experiments of r=0.85 (range 0.81-0.88) exceeds the minimum r≥0.8 standard defined by ENCODE for replicate WGBS experiments (**Fig. 1E**).

Of course, because an *in situ* Hi-C data set at loop resolution requires far more reads than a typical WGBS experiment (as reflected in, for instance, the ENCODE standards for the two protocols), the methylation track emerging from our loop resolution Hi-Culfite experiment in GM12878 has much deeper coverage than the corresponding ENCODE WGBS experiments (84X as compared to the ENCODE standard of 30X). It is notable that, despite the presence of a larger number of reads, the Hi-Culfite protocol described here covers slightly fewer CpG sites than WGBS: 88.3% versus 90.4% covered by at least one read (**Supplementary Fig. 1**). This is because CpGs far from a restriction site – typically those in low-complexity regions of the genome – are rarely covered using the Hi-Culfite protocol. However, when we examine CpGs that are covered multiple times in WGBS experiments, this effect disappears. For instance, 71.2% of CpGs are covered at least 10-fold in our Hi-Culfite experiment, versus 67.9% for ENCODE WGBS; and 39.5% of CpGs are covered 30-fold or higher in our Hi-Culfite experiment, versus 4.8% for ENCODE WGBS.

When we repeated the above analyses in other cell lines, including IMR90 (lung myofibroblasts) and HAP1 (haploid chronic myelogenous leukemia), we obtained similar results (**Fig. 1D**, **Supplementary Fig. 1**). We also confirmed that Hi-Culfite can recapitulate the effects of well-characterized perturbations to nuclear architecture and DNA methylation. For instance, 5-azacytidine is known to inhibit DNA methylation^13^. When we performed Hi-Culfite on GM12878 and on HapI cells treated with 5-azacytidine, we found that methylation declined genome-wide, as expected (on average by 2-fold see **Supplementary Fig. 2, Supplementary Table 2**). We did not discern an impact on nuclear architecture (**Supplementary Fig. 2**), consistent with the notion that DNA methylation may be dispensable for higher-order chromatin organization^14^. Degradation of cohesin is known to lead to the disappearance of CTCF-mediated looping^15,16,17^. When we degraded cohesin in HCT-116 colorectal carcinoma cells (**Online Methods**) and assayed the results using Hi-Culfite, we found that chromatin loops disappear, as expected (**Supplementary Fig. 3**). By contrast, we observed no effect on WGBS (**Supplementary Fig. 3**). Note that we used shallower maps for these analyses, ranging from 42 M to 171 M read pairs.

**Figure 2.**
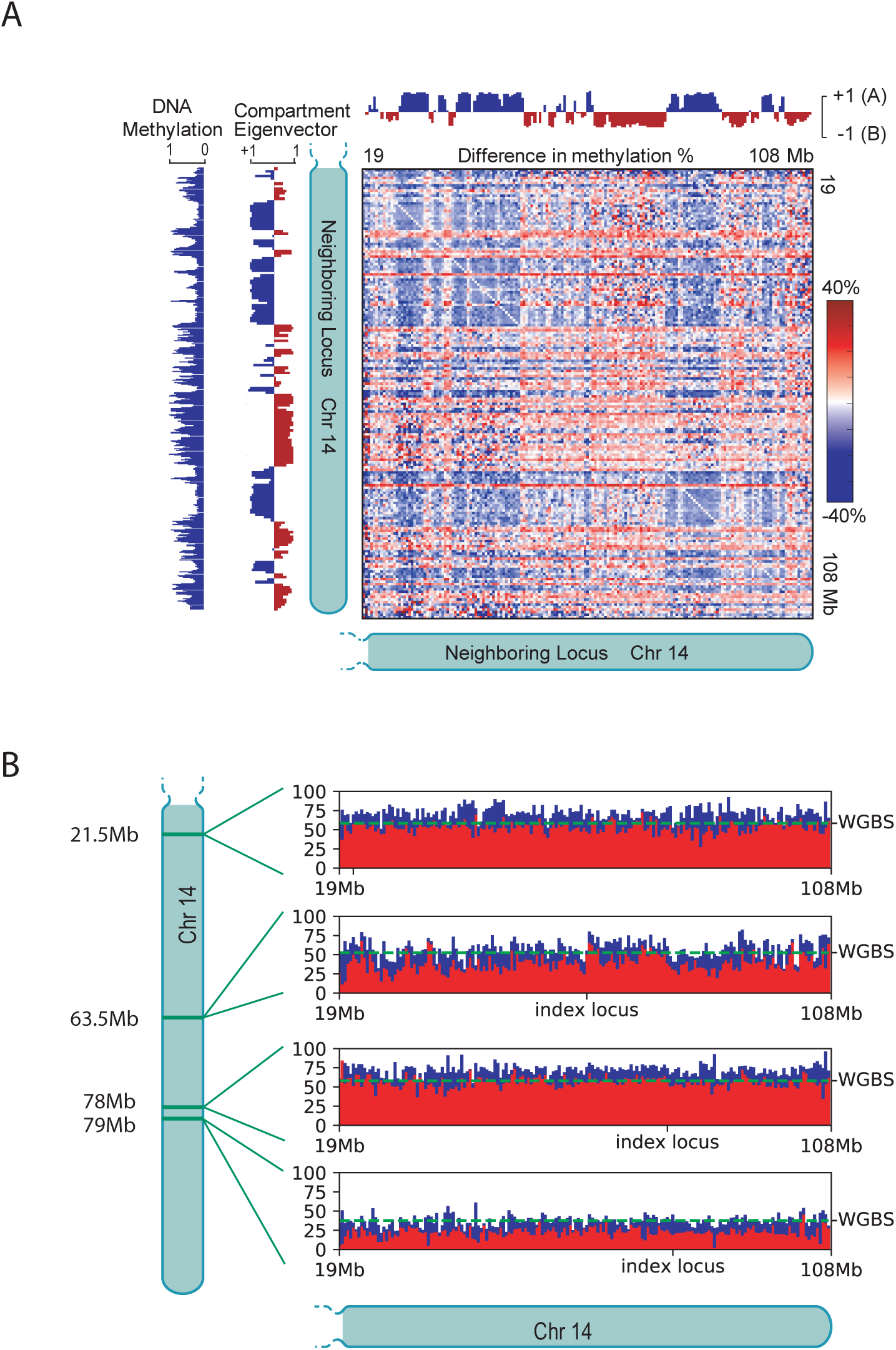
The methylation frequency of a locus varies as a function of its chromatin neighborhood in a particular cell. (**A**) The methylation frequency of a locus varies as a function of its chromatin neighborhood. Here, we show the deviation between the genome-wide methylation frequency for a 500-kb locus, and its observed methylation frequency when in contact with a particular neighbor locus. Elevated values are shown in red, and diminished values in blue. Loci in the A compartment are more likely to be methylated when in contact with a locus in the B compartment. (**B**) Methylation frequency as a function of the genomic position of the neighbor locus and its methylation state (**Online Methods**). When the neighbor locus is methylated, the methylation frequency at the index locus is indicated using the red bars; when the neighbor is unmethylated, the methylation frequency at the index locus is indicated using the blue bars. The dashed green line indicates the average methylation frequency at that locus.

Crucially, Hi-Culfite makes it possible not only to generate both the contact map and methylation profile data at once, but to perform integrative analysis of the underlying phenomena in ways that are not feasible when the assays are performed separately. In Hi-Culfite, each bisulfite-transformed read’s neighbor sequence provides additional information about its chromatin neighborhood. Since the vast majority (typically 75%) of ligations in an *in situ* Hi-C experiment happen in *cis*^8^, and nearly all of them happen in the same nucleus^8,18,19^, Hi-Culfite can provide insights about long-range epigenetic concordance and co-regulation that are not visible in ensemble DNA methylation measurements on heterogeneous cell populations.

For instance, to determine the effect of chromatin neighborhood on the methylation state of DNA, we partitioned the genome into loci of 500 kb each. We then calculated how often sequences derived from an index locus were methylated (i.e., exhibited mostly methylated CpGs) conditioned on the identity of the locus from which their neighbor sequence originated. Strikingly, we found that the methylation frequency of a given sequence was strongly associated with this spatial context. As an example, sequences deriving from the locus chr14: 37-37.5 Mb were methylated 64% of the time when the neighbor was locus chr14: 68.5-69 Mb (in 14 out of 22 cases), but only 6% of the time when the neighbor was locus chr14: 75.5-76 Mb (in 1 out of 17 cases).

We therefore generated a matrix showing the mean methylation frequency of every locus (i.e., what fraction of the time sequences from the locus were methylated) as a function of the identity of the neighboring locus. This revealed that the methylation frequency of reads derived from the A compartment depended especially strongly on their spatial context. When loci in the A compartment (open chromatin) had a neighbor in the B compartment (closed chromatin), they were methylated 34% more often than when their neighbor was in the A compartment. By contrast, loci in the B compartment exhibited less dependence on spatial context: when their neighbor was in the A compartment, they were methylated 7% less often than when their neighbor was in the B compartment (**Fig. 2A, Supplementary Figure 4**).

Next, we asked whether the methylation state of a read tended to correlate with the methylation state of the neighboring sequence. We found that, regardless of the identity of the neighboring locus or how far away it lay along the contour of the chromosome, the likelihood that a read was methylated was higher, increasing from 40% to 52% if the neighboring read was also methylated (**Fig. 2B**). This demonstrates that pairs of sequences that are in spatial proximity have correlated methylation states, regardless of how far apart those sequences lie in the genome. Finally, to facilitate multi-omics analysis of Hi-Culfite data by the scientific community, we enhanced the Juicer software for Hi-C data analysis^20^. Our new module, dubbed JuiceMe (http://github.com/aidenlab/juiceme), is designed to work with Hi-Culfite data, allowing users to create whole-genome contact maps, whole-genome bisulfite tracks, and quality control statistics relevant to both data types (**Fig. 1B; Online Methods** and **Supplementary Table 1**). In addition, we enhanced the Juicebox visualization system^21^ to enable interactive exploration of Hi-Culfite data (http://github.com/aidenlab/juicebox).

Taken together, our study introduces a new set of experimental and computational tools for simultaneously probing both the nucleome and the methylome. Our results demonstrate that the methylation state of a sequence is context-sensitive, depending not only on the loci in its spatial neighborhood, but also on their methylation state. More generally, our results highlight the ways in which Hi-Culfite, and other multi-omics approaches, enable us to integrate disparate data types into new insights about the interplay between cellular phenomena.

## Methods

See Online Methods.

## Supporting information

## Acknowledgements

Supported by an NIH New Innovator Award (1DP2OD008540), an NSF Physics Frontier Center Grant (PHY-1427654, Center for Theoretical Biological Physics), the USDA National Institute of Food and Agriculture (2017-05741), the Welch Foundation (Q-1866), an NVIDIA Research Center Award, an IBM University Challenge Award, a Google Research Award, a Cancer Prevention Research Institute of Texas Scholar Award (R1304), a McNair Medical Institute Scholar Award, an NIH Encyclopedia of DNA Elements Mapping Center Award (UM1HG009375), and the President’s Early Career Award in Science and Engineering (4DP2OD008540) to E.L.A., the Paul and Daisy Soros Fellowship for New Americans to M.S.S., and Paul and Daisy Soros Fellowship, a Fannie and John Hertz Foundation Fellowship, and a Cornelia de Lange Syndrome Foundation grant to S.S.P.R. We are grateful to Sheikh Russell and Ido Machol for help with experiments and the entire Aiden Lab for fruitful discussions. We would like to thank Sigrid Knemeyer for help with the figures. Thanks also to the ENCODE consortium labs of Myers, Bernstein, Snyder, Iyer, and Stamatoyannopoulos for the WGBS and CTCF data.

## Author Contributions

Author contributions are as follows: E.L.A. and S.S.P.R. conceived of this project; E.K.S. and O.D. designed the protocol; E.K.S., O.D., S-C.H. and I.D.B. performed experiments; N.C.D. built the computational pipeline; N.C.D., M.S.S., Y.J., and E.L.A. analyzed the data; E.K.S, N.C.D, M.S.S., E.S.L., A.G., and E.L.A. prepared the manuscript.

## Competing Interests

The authors declare no competing interests.

## Online Methods

### Hi-Culfite Protocol

#### Cell Culture

All commercial cell lines were cultured following vendors’ recommendations to about 80% confluency before they were harvested and cross-linked in fresh complete medium. For methylation inhibition experiments, cultures of GM12878 (Coriell Institute) and Hap1 (Horizon) were grown for 8 days in medium supplemented with DMSO without drug or with 1 μM or 5 μM 5-azacytidine (Sigma Aldrich) solubilized in DMSO, refreshing the medium/drug every two days.

IMR90 (CCL-186) cell line was purchased from ATCC and expanded as recommended.

We obtained HCT-116-CMV-OsTir1 and HCT-116-RAD21-mAID-mClover cells (HCT-116 RAD21-mAC) from Masato Kanemaki^24^. The cells were cultured in McCoy’s 5A medium supplemented with 10% FBS, 2 mM L-glutamine, 100 U/ml penicillin, and 100 μg/ml streptomycin at 37°C with 5% CO2. Degradation of the AID-tagged RAD21 was induced by the addition of 500 μM indole-3-acetic acid (IAA; Sigma Aldrich). For experiments with untreated cells and cells treated for 6 hours, medium was aspirated at t=0, and either replaced with fresh medium (untreated) or medium containing 500 μM IAA. The cells were then washed, trypsinized and processed for downstream experiments at t=6hrs.

#### Library Construction

Hi-C libraries were prepared using the protocol described in^8^ Briefly, one million cells were crosslinked with 1% formaldehyde for 10 minutes at room temperature and then quenched with 0.2 M glycine solution. Cells were lysed and nuclei permeabilized with 0.5% SDS for 10 minutes at 62°C. Chromatin was digested with 100 U of MboI restriction enzyme (NEB). Ends of the restriction fragments were filled-in and labeled with a biotinylated nucleotide and then ligated. Nuclei were pelleted, proteins were digested with proteinase K and crosslinks were reversed by heating at 68°C overnight. DNA was sheared in a Covaris focused ultrasonicator to a length of 300-500 bp. Size-selected DNA was split for processing with two workflows – 10% of the material was used for preparation of a regular Hi-C library (unconverted control) and 90% of the DNA was used for Hi-Culfite library construction. Hi-C libraries were finished by enriching for biotinylated ligation junctions through binding to T1 streptavidin beads (Thermo Fisher) and preparing the library for Illumina sequencing performing the end-repair, A-tailing and adapter ligation steps with DNA attached to the beads. Libraries were amplified directly off the beads and purified for subsequent Illumina sequencing.

DNA for Hi-Culfite (see Protocol Exchange for detailed step-by-step protocol) was first treated with sodium bisulfite using EpiTect Fast bisulfite conversion kit (Qiagen) following the kit’s instructions and extending each of the two 60°C conversion incubations to 20 minutes. Converted DNA was purified without addition of an RNA carrier to the binding buffer. Biotinylated ligation junctions in purified bisulfite-converted DNA were captured on 15 μl C1 streptavidin beads (Thermo Fisher) in denaturing binding buffer (10 mM Tris pH 7.5; 5 mM EDTA; 500 mM LiCl; 0.5% Igepal CA630; 0.2% SDS; 4 M Urea) for 10 minutes at 55°C. DNA was washed twice with binding buffer at 55°C and once with 10mM Tris. Beads were resuspended in 15 μl of 10 mM Tris and libraries were detached from the beads by heating for 5 minutes at 95°C. After separating on a magnet, library construction for Illumina sequencing was performed on the supernatant containing ssDNA library using Accel-NGS Methyl-Seq kit (Swift Biosciences) following the kit’s manual. Libraries were amplified with barcoded primers using 8-10 amplification cycles. Final libraries were purified and molecules in the range of 450-650 bp were selected by agarose gel electrophoresis and subsequent gel extraction.

Both Hi-C and Hi-Culfite libraries were first sequenced with 80 bp paired-end reads on an Illumina NextSeq instrument, obtaining about 2 million read pairs per library. Data quality was evaluated and successful libraries were then deep sequenced with 150 bp paired-end reads on the Illumina HiseqX platform.

### JuiceME: Hi-Culfite Data Processing Pipeline

The data processing pipeline for Hi-Culfite is a modified version of the Juicer pipeline^20^. Since the DNA has been bisulfite-converted, the aligner must be able to handle mapping to essentially two different genomes. Additionally, after alignment the reads must be combined to generate WGBS sequencing tracks. The other steps of the Juicer pipeline (chimera handling, duplicate removal, Hi-C contact map creation and normalization) remain the same.

#### Sequence Alignment with bwa-meth

All Hi-Culfite data reported in this paper was generated using Illumina paired-end sequencing. The sequencer produces two fastq files, one for each read end. As with any proximity ligation assay, each read end must be aligned separately as a single end read so that the aligner does not make incorrect assumptions about the insert size.

We use *bwa-meth*^22^ as our aligner. *Bwa-meth* uses *bwa*^*25*^ as its base aligner, which performs well on Hi-C data^20^. *Bwa-meth* works by creating an alternate methylated version of the genome and then calling *bwa* to align. Since each read end is aligned separately, we must first reverse complement the second read end before calling *bwa-meth*.

All reads that align are merged into a BAM file that is coordinate-sorted for methylation processing. For the Hi-C contact map creation, the rest of the pipeline proceeds exactly as previously described^20^: chimeras are appropriately handled, duplicates and near duplicates are removed, and contact maps are created and normalized.

#### Methylation Track Generation with MethylDackel

Duplicate Hi-C contacts are marked as duplicates in the methylation BAM. We then call the program *MethylDackel extract* with the flag “-F 1024”; this ignores duplicates reads but keeps all other mapped reads with MAPQ ≥ 10. *MethylDackel* generates CpG methylation tracks, a cytosine coverage report, and an input file for the analysis program *MethylKit*^26^. We used *MethylKit* to produce the correlation analysis in **Fig. 1**. The CpG methylation tracks are bedGraph; we used *igvtools*^27^ to create a TDF file for fast viewing.

We call *MethylDackel perRead* to produce methylation information per read instead of the usual per cytosine. The results from this program are then combined with the list of Hi-C contacts in order to create binned contact maps separated by methylation status. Each contact that has methylation status information on both read ends is classified as either “both methylated” (both read ends are methylated), “both unmethylated” (both read ends are unmethylated), or “methylated-unmethylated” (one read end is methylated and the other is unmethylated). These contact maps are used in the co-methylation analysis, described below.

### Neighborhood Methylation Analysis

For the neighborhood methylation analysis, we would like to determine how the chromatin neighborhood (i.e., the loci that any given locus is in contact with) affects the methylation state of the DNA of that locus. That is, we want to know the methylation percentage of locus *i* given that it interacts with locus *j*.

Each Hi-C contact in this analysis has a methylation status of 0 or 1 on each read end, based on whether or not the methylation status of the CpGs it covers result in >50% methylation. We can then split Hi-C contacts into four different matrices: contacts in which both read ends are methylated, contacts in which both read ends are unmethylated, contacts in which read end *i* is methylated and read end *j* is unmethylated, and contacts in which read end *i* is unmethylated and read end *j* is methylated. The latter two matrices are transposes of one another, so we define these matrices as, respectively, *M*, *U*, *Y*, and *Y*^*T*^.

Then the probability that locus *i* is methylated given that it is in contact with locus *j* is the sum of contacts at locus *i,j* in which locus *i* is methylated, divided by the total number of contacts at locus *i,j*.

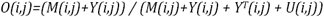

Now consider the null hypothesis. If there were no effect of neighborhood on methylation status, we would expect the methylation percentage of locus *i* given that it interacts with locus *j* to be the same regardless of locus *j*. Define *a* as the one-dimensional average methylation at locus *i*:

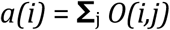

Then

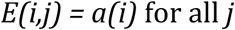

The matrix in **Fig. 2A** is the *O* minus *E*; its entries give a measure of methylation frequency divergence from the expected model. Where *O-E* is 0, the data is compatible with the null model, i.e. solely based on the overall average methylation at that locus. High or low values indicate a divergence whereby locus *i* is more or less methylated than we would expect due to its interaction with locus *j*.

### Methylation Correlation Analysis

For the methylation correlation analysis, we would like to determine if the methylation state of a read correlates with the methylation state of the neighboring sequence. We define the methylation correlation as the frequency with which locus *j* is methylated given that locus *i* is methylated, divided by the total number of times locus *j* is methylated. This is

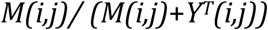

Similarly, we examine the unmethylation correlation: the number of times *j* is unmethylated given that *i* is unmethylated, divided by the total number of times locus *j* is unmethylated. This is

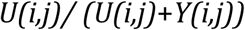

If methylation status were uncorrelated with the methylation state of the neighboring sequence, rows of these correlation matrices would simply equal the average methylation status at locus *i*, i.e. *a(i)*. As we show in **Fig. 2B**, this is not the case; the methylation likelihood changes when the neighboring read is methylated.

### Supplemental Co-methylation Analysis

For the comethylation analysis, we want to determine if Hi-C contacts have both ends methylated or both ends unmethylated at a higher frequency than one would expect given the baseline methylation of the loci. Using the matrices defined above, the observed comethylation frequency is

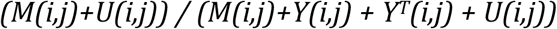

The expected comethylation frequency is calculated from the average methylation vector *a*. It is the probability that both are methylated plus the probability that both are unmethylated:

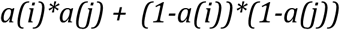

In **Supplemental Figure 4**, we show the observed comethylation frequency minus the expected comethylation frequency. The matrix is mostly positive, indicating that loci are comethylated more often than the null hypothesis would indicate.

### Supplemental Aggregate Methylation Analysis

GM12878 CTCF peak calls from ENCODE^11,12^ were intersected to find common CTCF peaks. This set of peaks was intersected with the HG19 CTCF motif database hosted for Juicer^20^, which was originally built using FIMO^28^. CTCF motifs peaks were split into forward and reverse motifs. Forward and reverse CTCF motif peaks were further subdivided into looping and non-looping motifs, by their presence or absence in the GM12878 loop list with motifs^8^.

Methylation data was generated using the JuiceMe pipeline for the respective Hi-C experiments. Bedgraph files were converted to bigwig files using UCSC executables^29^. Aggregation analysis using the CTCF motif peaks and methylation data was performed using bwtool^30^, and post-processed and visualized using python code hosted at github.com/aidenlab/JuiceMe.

### Supplemental Aggregate Peak Analysis (APA)

APA was performed using Juicer Tools^3^ on the Hi-C maps at 25kb (unless otherwise specified), using loop lists for the maps generated from prior experiments. Loop lists for GM12878, HapI, and HCT-116 were previously published^8,15,31^.

