## Supplementary material for "The Hi-Culfite assay reveals relationships between chromatin contacts and DNA methylation state"

### Supplemental Materials

### Aggregate Methylation Analysis

In some cases, subtle effects in WGBS data are revealed only when studied in aggregate (i.e., summing the methylation profile at a large number of positions), but are not clearly visible when loci are examined one at a time. Having confirmed that the methylation profiles produced by Hi-Culfite data match WGBS-derived profiles across loci and resolutions, we wondered whether Hi-Culfite data would recapitulate these aggregate effects as well. For example, aggregate analyses of WGBS data have shown that DNA methylation is depleted at CTCF-bound sites[1] and negatively correlated with nucleosome occupancy in the flanking regions[2,3]. To confirm that this relationship is accurately observed using Hi-Culfite, we plotted DNA methylation in the vicinity of CTCF motifs and compared it to nucleosome occupancy data from MNase-Seq[4,5] (DCC accession ENCSR000CXP). Indeed, we observed the DNA methylation is greatly depleted at CTCF-bound sites, and that the flanking regions exhibit peaks and troughs in methylation that are exactly out of phase with the position of nucleosomes (**Supplementary Fig. 5**). Moreover, it is well-known that loops form between CTCF-bound anchor sites in the convergent orientation, i.e., with the motifs pointing towards one another[6]. Comparing DNA methylation on opposite sides of a CTCF-bound motif, we found that the DNA methylation level is consistently diminished on one side, corresponding to the interior of the loop (**Supplementary Fig. 5**). This confirms an observation first reported in[3]. These results further confirm the ability of Hi-Culfite to detect subtle effects that can only be observed when methylation profiles at many loci are studied in aggregate.

**Supplementary Figure 1. Comparison of Hi-Culfite data to separate WGBS and Hi-C data sets.** (A) CpG read coverage percentage in GM12878 WGBS data from two replicates generated by ENCODE Consortium and two replicates of Hi-Culfite of GM12878.
(B) Low sequencing depth comparison of contact maps, DNA methylation tracks (blue) and eigenvectors indicating open (positive values) and closed (negative values) chromatin compartments along chromosome 14 of Hap1 cell line generated by single Hi-Culfite experiment (left) and separate in situ Hi-C and WGBS assays (right). (C) Scatter plot style heat map (bottom left), Pearson correlation coefficients for pair-wise comparisons (top right) and bimodal distributions of single-CpG DNA methylation values (diagonal) for HapI and IMR90. Hi-Culfite data generated for both cell lines was compared with respect to WGBS was from HapI and ENCODE WGBS data for IMR90. This plot was produced by *MethylKit*[7].

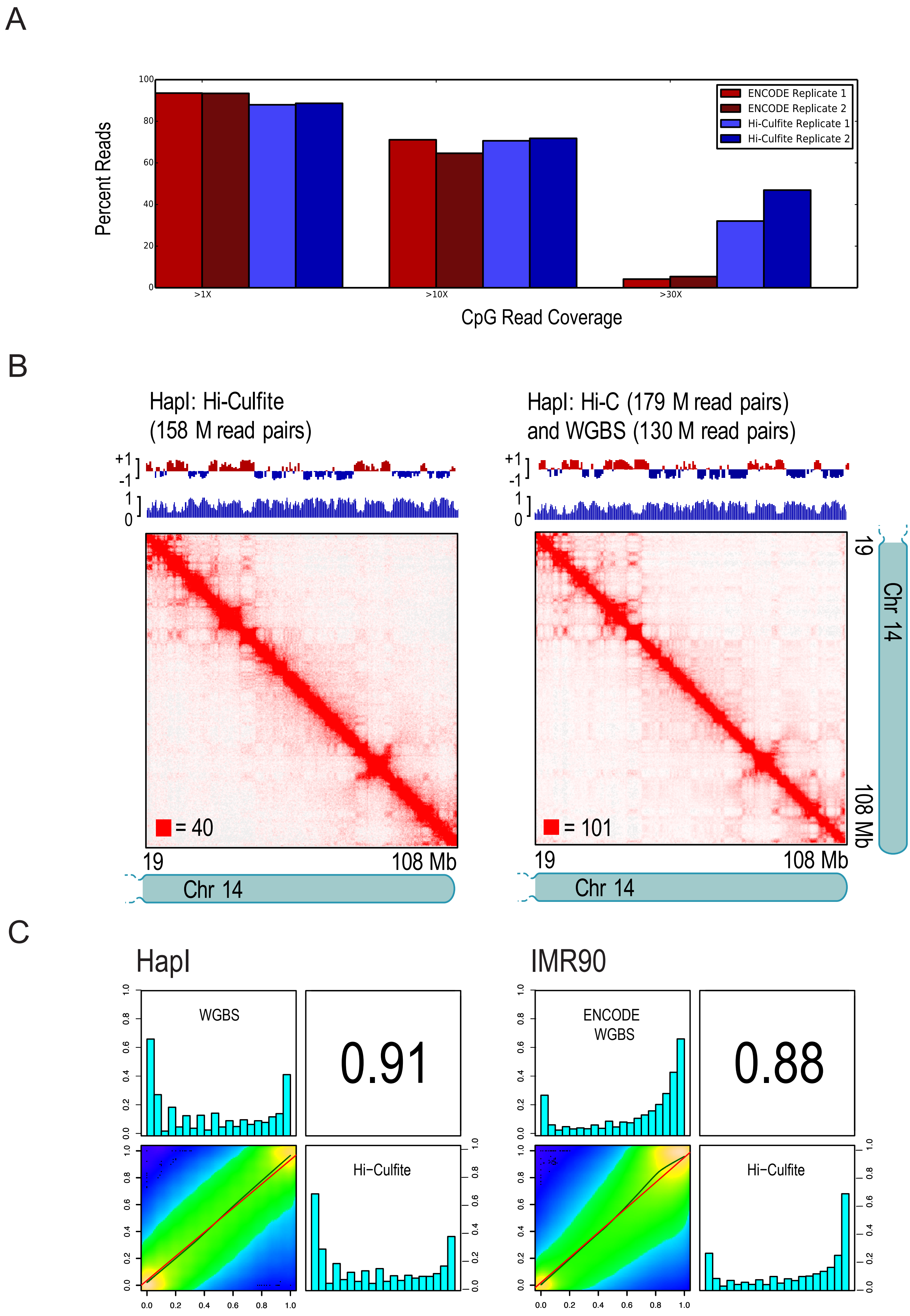

A
Percent Reads
ENCODE Replicate 1
ENCODE Replicate 2
Hi-Culfite Replicate 1
Hi-Culfite Replicate 2
>1X
>10X
>30X
CpG Read Coverage
B
HapI: Hi-Culfite
(158 M read pairs)
HapI: Hi-C (179 M read pairs)
and WGBS (130 M read pairs)
= 40
= 101
19
108 Mb
Chr 14
C
HapI
IMR90
WGBS
0.91
Hi−Culfite
ENCODE
WGBS
0.88
Hi−Culfite

**Supplementary Figure 2. Hi-Culfite data shows that DNA methylation inhibition by 5-azacytidine has no effect on chromosome-level nuclear organization in GM12878 and Hap1 cell lines.**

(A) Hi-Culfite contact maps of chromosome 14 and eigenvectors indicating open (positive values) and closed (negative values) chromatin compartments of GM12878 and Hap1 cells treated for 8 days with DMSO (control) or increasing concentration (1uM and 5uM) of 5-azacytidine show no difference despite of global decrease in DNA methylation (methylation tracks in blue) (B) Aggregate Peak Analysis performed on the Hi-C maps using loop lists for GM12878 and HapI show no significant changes in looping signal with increasing 5-azacytidine concentration. In order to assess the potential formation of new convergent CTCF loops with 5-azacytidine treatment, loop lists from the other cell type were also used for APA analysis on the alternative map.

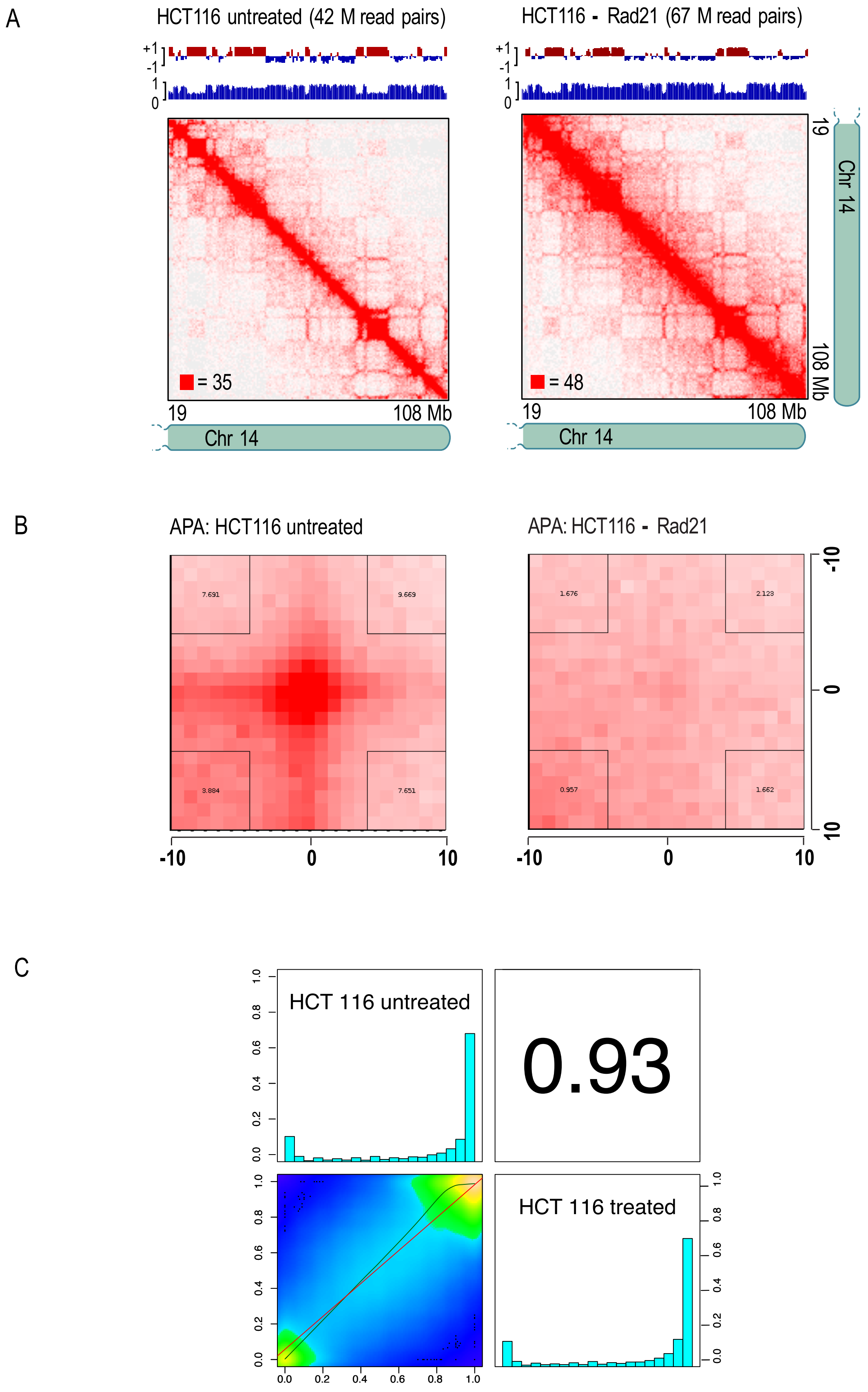

A
HCT116 untreated (42 M read pairs)
HCT116 - Rad21 (67 M read pairs)
+1
-1
1
0
= 35
= 48
19
108 Mb
Chr 14
B
APA: HCT116 untreated
APA: HCT116 - Rad21
7.691
9.669
3.884
7.651
1.676
2.123
0.957
1.662
-10
0
10
C
HCT 116 untreated
0.93
HCT 116 treated
0.0
0.2
0.4
0.6
0.8
1.0

**Supplementary Figure 3. RAD21 Degron Hi-Culfite experiments recapitulate loss of CTCF-mediated loops.** (A) Hi-Culfite contact maps of chromosome 14 and eigenvectors indicating open (positive values) and closed (negative values) chromatin compartments of HCT-116 before and after auxin treatment. Aggregate Peak Analysis performed at 10 kB resolution using the HCT-116 loop list shows loss of loops with auxin treatment.

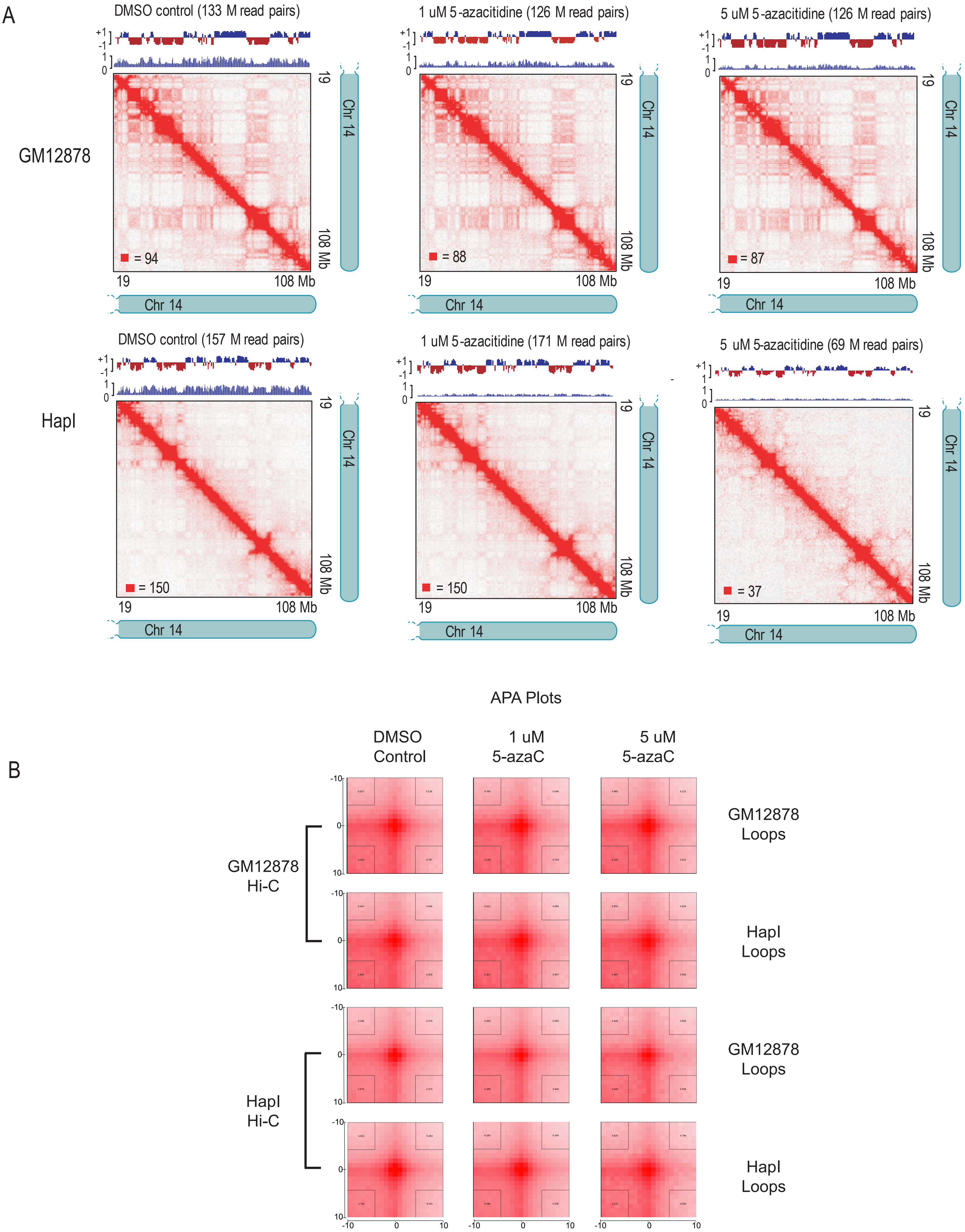

A
DMSO control (133 M read pairs)
1 uM 5-azacitidine (126 M read pairs)
5 uM 5-azacitidine (126 M read pairs)
GM12878
= 94
= 88
= 87
DMSO control (157 M read pairs)
1 uM 5-azacitidine (171 M read pairs)
5 uM 5-azacitidine (69 M read pairs)
HapI
= 150
= 150
= 37
19
108 Mb
Chr 14
APA Plots
B
DMSO Control
1 uM 5-azaC
5 uM 5-azaC
GM12878 Hi-C
HapI Hi-C
GM12878 Loops
HapI Loops
GM12878 Loops
HapI Loops

**Supplementary Figure 4. Comethylation analysis shows that Hi-C contacts share methylation status more frequently than expected from the null hypothesis.** (A) Difference of observed comethylation matrix and expected comethylation matrix on chromosome q14 at 1Mb. The difference is overwhelmingly red, indicating that Hi-C contacts tend to share the same methylation status. (B) Difference of the observed methylation correlation matrix and the observed unmethylation correlation matrix. A methylated locus is more likely to depend on its spatial context than an unmethylated locus.

A

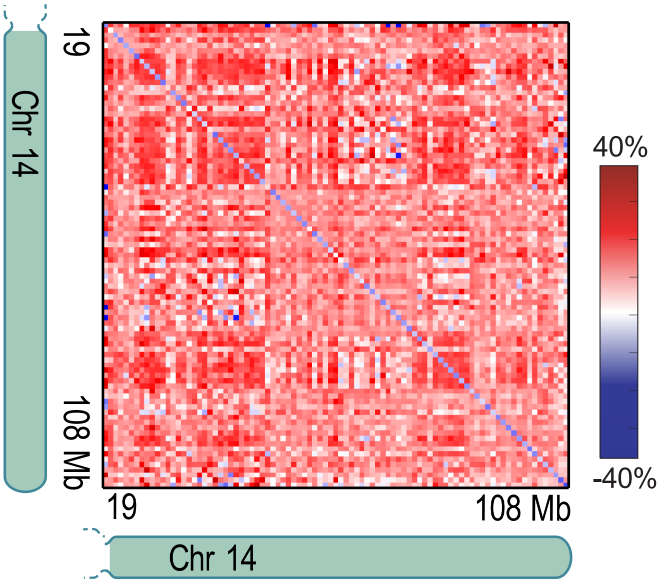
19
108 Mb
Chr 14
19
108 Mb
Chr 14
40%
-40%

B

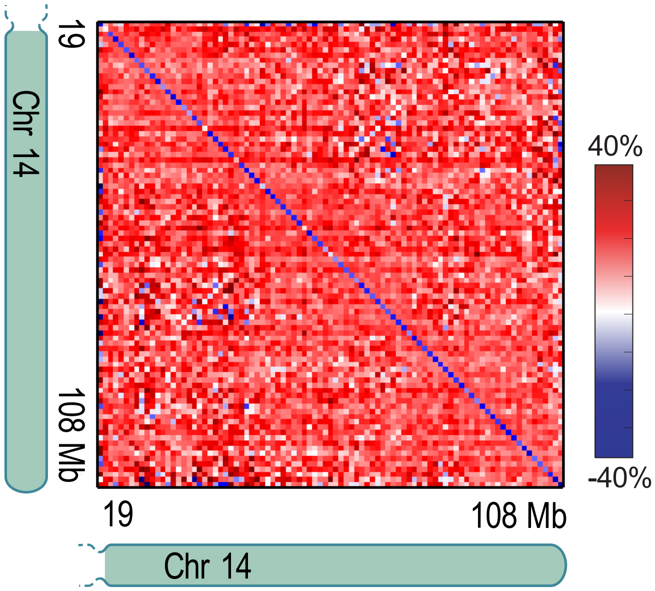
19
108 Mb
Chr 14
19
108 Mb
Chr 14
40%
-40%

**Supplementary Figure 5. Aggregate methylation analysis recapitulates CTCF motif site relationships with nucleosome occupancy and methylation asymmetries.** (A) Aggregation of methylation data from the combined Hi-Culfite GM12878 replicates 1 and 2 at oriented CTCF motifs recapitulates nucleosome occupancy and directional methylation asymmetry. (B) Aggregation of methylation data from 5-azacytidine experiments shows no changes to nucleosome occupancy and directional methylation asymmetry with increasing concentration of 5-azacytidine.

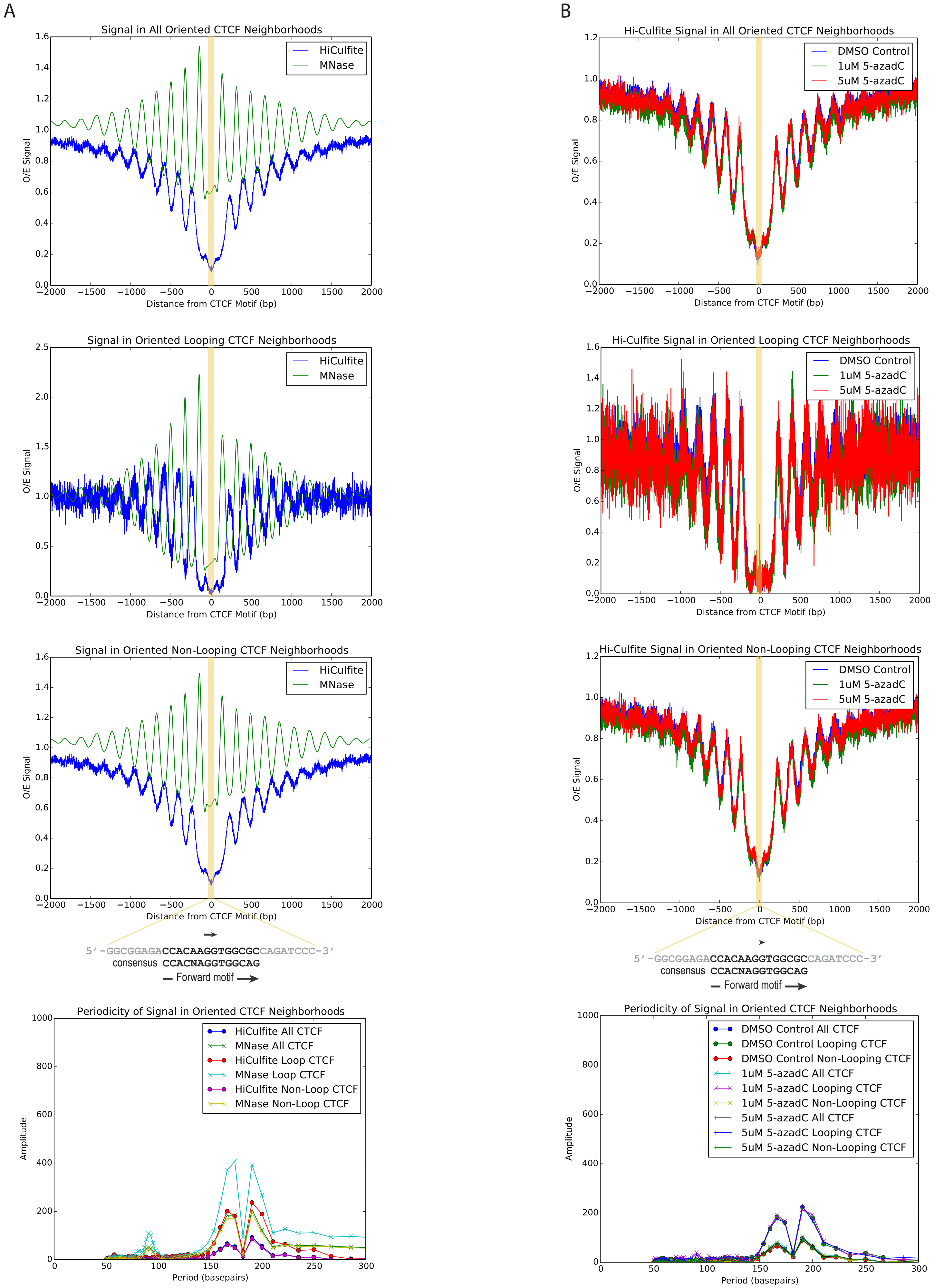
A
Signal in All Oriented CTCF Neighborhoods
HiCulfite
MNase
O/E Signal
Distance from CTCF Motif (bp)
Signal in Oriented Looping CTCF Neighborhoods
Signal in Oriented Non-Looping CTCF Neighborhoods
5′-GGCGGAGACCACAAGGTGGCGCCAGATCCC-3′
consensus CCACNAGGTGGCAG
Forward motif
Periodicity of Signal in Oriented CTCF Neighborhoods
HiCulfite All CTCF
MNase All CTCF
HiCulfite Loop CTCF
MNase Loop CTCF
HiCulfite Non-Loop CTCF
MNase Non-Loop CTCF
Amplitude
Period (basepairs)
B
Hi-Culfite Signal in All Oriented CTCF Neighborhoods
DMSO Control
1uM 5-azadC
5uM 5-azadC
Hi-Culfite Signal in Oriented Looping CTCF Neighborhoods
Hi-Culfite Signal in Oriented Non-Looping CTCF Neighborhoods
DMSO Control All CTCF
DMSO Control Looping CTCF
DMSO Control Non-Looping CTCF
1uM 5-azadC All CTCF
1uM 5-azadC Looping CTCF
1uM 5-azadC Non-Looping CTCF
5uM 5-azadC All CTCF
5uM 5-azadC Looping CTCF
5uM 5-azadC Non-Looping CTCF

| | Hi-Culfite GM12878 | Hi-C GM12878 |
|---|---|---|
| **Sequenced Read Pairs:** | 1,748,191,006 | 2,935,830,058 |
| **Alignment (% Sequenced Reads)** | | |
| **Normal Paired:** | 1,092,961,004 (62.52%) | 2,187,069,164 (74.50%) |
| **Chimeric Paired:** | 518,160,223 (29.64%) | 507,602,762 (17.29%) |
| **Chimeric Ambiguous:** | 111,331,581 (6.37%) | 110,359,127 (3.76%) |
| **Unmapped:** | 25,738,198 (1.47%) | 130,799,005 (4.46%) |
| **Duplication and Complexity (% Sequenced Reads)** | | |
| **Alignable (Normal+Chimeric Paired):** | 1,611,121,227 | 2,694,671,926 |
| **Unique Reads:** | 1,336,677,373 | 2,584,147,879 |
| **PCR Duplicates:** | 268,384,692 | 110,448,756 |
| **Optical Duplicates:** | 6,059,162 | 75,291 |
| **Analysis of Unique Reads (% Sequenced Reads / % Unique Reads)** | | |
| **Intra-fragment Reads:** | 149,415,012 (8.55% / 12.18%) | 77,475,958 (2.64% / 3.00%) |
| **Below MAPQ Threshold:** | 186,326,300 (10.66% / 15.19%) | 238,800,841 (8.13% / 9.24%) |
| **Hi-C Contacts:** | 890,634,698 (50.95% / 72.62%) | 2,267,880,754 (77.25% / 87.76%) |
| **3' Bias (Long Range):** | 56% - 44% | 71% - 29% |
| **Pair Type % (L-I-O-R):** | 25% - 25% - 25% - 25% | 25% - 25% - 25% - 25% |
| **Analysis of Hi-C Contacts (% Sequenced Reads / % Unique Reads)** | | |
| **Inter-chromosomal:** | 247,760,751 (14.17% / 20.20%) | 577,717,760 (19.68% / 22.36%) |
| **Intra-chromosomal:** | 642,873,947 (36.77% / 52.42%) | 1,690,162,994 (57.57% / 65.41%) |
| **Short Range (<20Kb):** | 200,014,362 (11.44% / 16.31%) | 539,732,380 (18.38% / 20.89%) |
| **Long Range (>20Kb):** | 442,758,907 (25.33% / 36.10%) | 1,150,429,110 (39.19% / 44.52%) |

**Table S1**
Quality metrics of Hi-Culfite libraries (two combined replicates) compared to in situ Hi-C (eight combined replicates, Rao et al. Cell 2014)

| GM12878 treated with 5-azacytidine | | | |
|---|---|---|---|
| | **Control** | **1 μM** | **5 μM** |
| **C->T conversion** | 99.41% | 99.10% | 99.46% |
| **CHH+CHG coverage** | 73.84% | 73.11% | 72.91% |
| **CpG C->T conversion** | 45.37% | 69.46% | 75.08% |
| **CpG coverage** | 71.59% | 70.71% | 70.30% |
| **Hap-1 treated with 5-azacytidine** | | | |
| | **Control** | **1 μM** | **5 μM** |
| **C->T conversion** | 99.06% | 99.37% | 98.82% |
| **CHH+CHG coverage** | 40.51% | 67.37% | 33.00% |
| **CpG C->T conversion** | 37.66% | 86.16% | 85.79% |
| **CpG coverage** | 37.60% | 65.22% | 30.99% |

**Table S2**
Methylation analysis metrics for Hi-Culfite libraries prepared from GM12878 and Hap1 cells treated for 8 days with DMSO (control), 1 μM or 5 μM 5-azacytidine in DMSO.
